# Molecular recognition of an aversive odorant by the murine trace amine-associated receptor TAAR7f

**DOI:** 10.1101/2023.07.07.547762

**Authors:** Anastasiia Gusach, Yang Lee, Armin Nikpour Khoshgrudi, Elizaveta Mukhaleva, Ning Ma, Eline J. Koers, Qingchao Chen, Patricia C. Edwards, Fanglu Huang, Jonathan Kim, Filippo Mancia, Dmitry B. Verprintsev, Nagarajan Vaidehi, Simone N. Weyand, Christopher G. Tate

## Abstract

There are two main families of G protein-coupled receptors that detect odours in humans, the odorant receptors (ORs) and the trace amine-associated receptors (TAARs). Their amino acid sequences are distinct, with the TAARs being most similar to the aminergic receptors such as those activated by adrenaline, serotonin and histamine. To elucidate the structural determinants of ligand recognition by TAARs, we have determined the cryo-EM structure of a murine receptor, mTAAR7f, coupled to the heterotrimeric G protein G_s_ and bound to the odorant N,N-dimethylcyclohexylamine (DMCH) to an overall resolution of 2.9 Å. DMCH is bound in a hydrophobic orthosteric binding site primarily through van der Waals interactions and a strong charge-charge interaction between the tertiary amine of the ligand and an aspartic acid residue. This site is distinct and non-overlapping with the binding site for the odorant propionate in the odorant receptor OR51E2. The structure, in combination with mutagenesis data and molecular dynamics simulations suggests that the activation of the receptor follows a similar pathway to that of the β-adrenoceptors, with the significant difference that DMCH interacts directly with one of the main activation microswitch residues.

## Introduction

Perception and interpretation of odours are essential for the life of vertebrates. Odorant molecules are detected in the nasal cavity by G protein-coupled receptors (GPCRs) in olfactory sensory neurons, which then transmit a signal to the olfactory bulb in the brain^1^. Each olfactory sensory neuron specifically expresses a single chemosensory GPCR that is activated by one or several volatile odorant molecules^2–5^. Detection of thousands of different odours is then possible through the action of hundreds of receptor types in their corresponding neurons. Approximately half of the ∼800 human GPCRs are chemosensory receptors^6^. These belong to the rhodopsin-like class A family and can be divided into two groups, the odorant receptors (ORs) and the trace amine-associated receptors (TAARs). Although rhodopsin-like GPCRs are the most abundant receptors in humans and the most well-studied from the structural and functional perspectives^7,8^, only recently has the first structure of an OR been determined^9^.

The TAARs are a small family of specialised receptors with only 6 representatives encoded by the human genome, compared to about 400 OR genes. The amino acid sequence of the TAAR studied here, TAAR7f, is most similar to aminergic receptors, specifically the β_2_-adrenoceptor (β_2_AR), rather than to ORs (Fig. 1a). TAARs bind trace amines^3,4^ that are typically small volatile molecules formed by the decarboxylation of amino acids^10^. These molecules serve as sensory cues for a range of stimuli, such as the presence of predators or prey, the proximity of a mating partner and the spoilage of food^10^, and elicit either attraction or aversion responses, depending on the odour. TAAR7f is a well-studied murine homologue of human TAAR9 (sequences are 72 % identical) with well-characterised agonists^11,12^. Mice elicit either attractive/neutral or aversive behaviour when exposed to TAAR7f ligands, such as amines found in urine^13^, with the physiological response dependent upon the ligand and its concentration^11^.

**Fig. 1.**
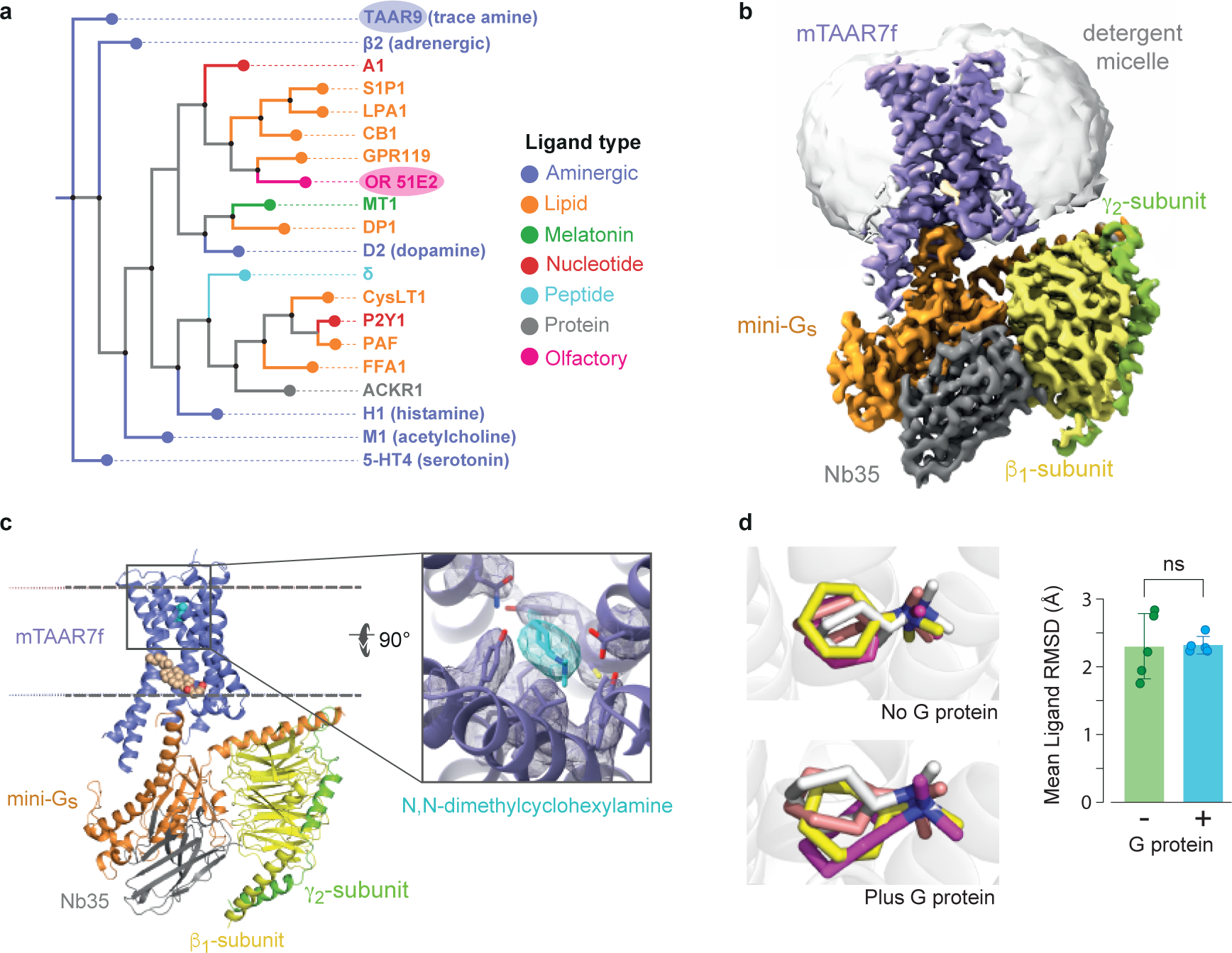
Phylogenetic analysis of selected human GPCRs and overall structure of the mTAAR7f-Gs complex. **a**, Representative human receptors of GPCR families interacting with different ligand types are compared to hTAAR9, the closest human homologue of mTAAR7f (72% amino acid residue identity). The phylogenetic analysis tool from GPCRdb.org was used. **b**, Cryo-EM density of the entire complex. **c**, cartoon of the mTAAR7f-Gs complex (ribbon representation) with bound DMCH (pale blue) and CHS (pale brown) shown as spheres. The inset shows density for DMCH (pale blue) and surrounding residues (purple) in mesh. The view is from the extracellular surface and is 90° orthogonal to the receptor cartoon viewed in the membrane plane. **d**, The three most populated ligand binding poses derived from MD simulations conducted either in the absence or presence of the G protein (ligand orientation from the cryo-EM structure is shown in grey). The bar graph shows differences in ligand RMSD from five distinct MD simulations either with or without G protein. The error bars represent the SD and a t-test showed no statistical difference (ns) between the mean ligand RMSDs.

### Structure determination of TAAR7f

In order to define the molecular recognition determinants of odorants by TAAR receptors, we determined a structure of murine TAAR7f (mTAAR7f) by electron cryo-microscopy (cryo-EM) in an active state coupled to the heterotrimeric G protein, G_s_. mTAAR7f was chosen from a screen of multiple different olfactory receptors as being a highly expressed receptor in insect cells using the baculovirus expression system and also being relatively stable after detergent solubilisation as assessed by fluorescence-detection size exclusion chromatography (FSEC). In addition, TAAR7f is known to bind the agonist N,N-dimethylcyclohexylamine (DMCH) with an EC_50_ of 0.5 μM^12^.

Wild-type mTAAR7f was tagged at the N-terminus (haemagglutinin signal sequence, FLAG tag, His_10_ purification tag and tobacco etch virus cleavage site) and C-terminus (human rhinovirus 3C cleavage site and eGFP). The construct was expressed in insect cells using the baculovirus expression system and purified in the presence of the agonist DMCH (see Methods; Extended Data Figs. 1 and 2a,b). *In vivo*, mTAAR7f couples to the heterotrimeric G protein G_olf_^14^. However, G_olf_ and G_s_ have very similar amino acid sequences (77% identical) and the molecular determinants of G_s_ coupling identified in GPCR-G_s_ structures^15^ were predicted to be identical in G_olf_, assuming that the respective G proteins couple in a similar fashion. We therefore used G_s_ for making a mTAAR7f complex, because of the availability of nanobody Nb35 that stabilises the interface between the α-subunit and β-subunit of the heterotrimeric G protein; this may have proved important in improving the stability of the complex^16^.

**Figure 2.**
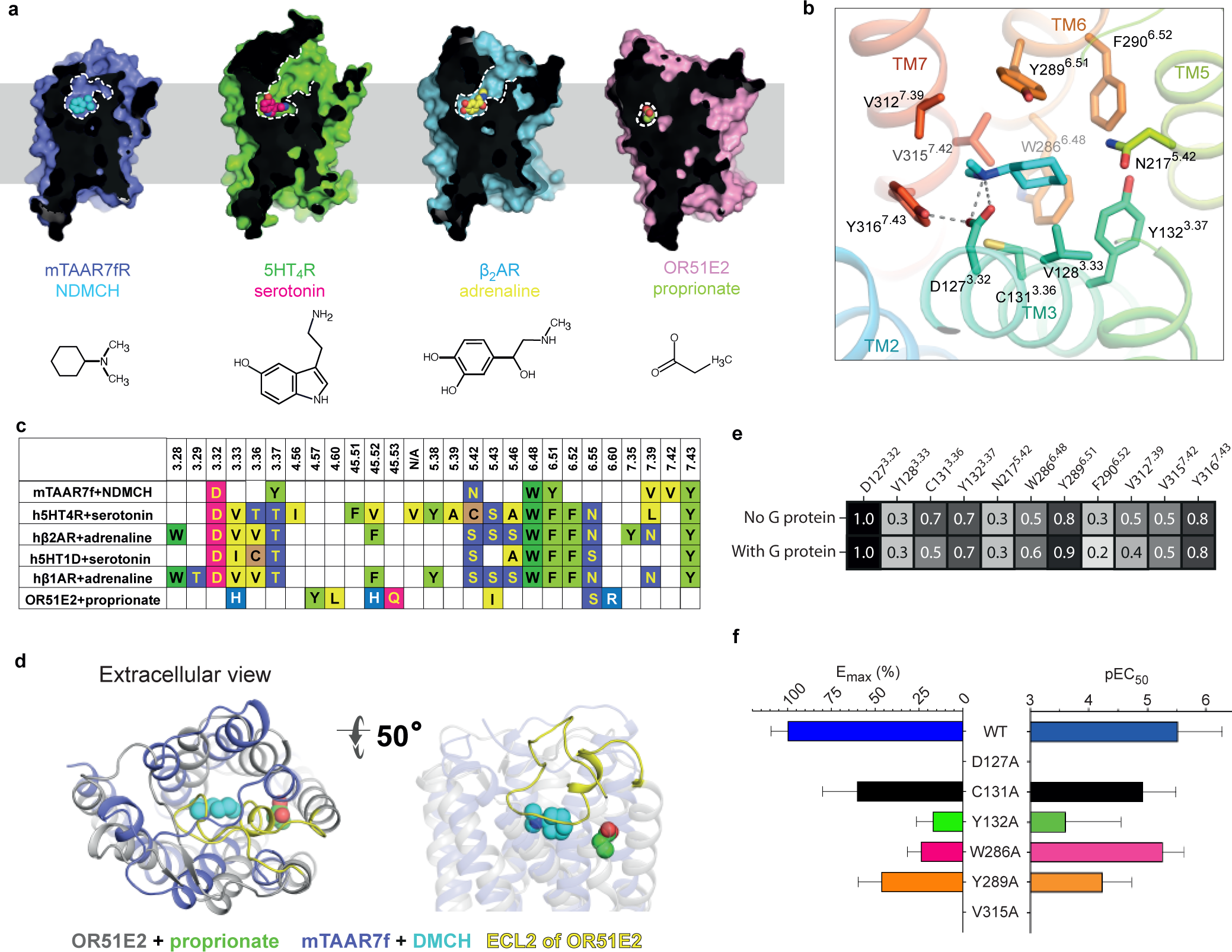
The mTAAR7f orthosteric binding site and comparison to other receptors. **a**, Sliced surface representation of the OBS of DMCH-bound mTAAR7f, serotonin-bound 5HT_4_R, adrenaline-bound β_2_AR and propionate-bound OR51E2; ligand atoms are depicted as spheres and the structures are shown below. **b,** Binding pose of DMCH and details of ligand-receptor interactions. Amino acid residues ≤ 3.9 Å from the ligand are shown with polar interactions depicted as dashed lines. **c,** Amino acid residues in the OBS within 3.9 Å of the ligand of mTAAR7f, aminergic receptors and an odorant receptor, OR51E2: (PDB IDs; h5HT_4_R, 7XT8; β_1_AR, 7JJO; β_2_AR, 4LDO; h5HT_1D_R, 7E32; OR51E2, 8F76). Numbers refer to the Ballesteros-Weinstein naming convention^23^. **d,** The relative positions of the OBS in OR51E2 and mTAAR7f are shown after superposition of the receptors. Ligands (propionate and DMCH) are shown as spheres. **e**, Frequency of ligand contacts as determined during MD simulations. **f**, G protein recruitment was assayed using BRET arising from NanoLuc-labelled receptor and Venus-labelled mini-G_s_. The mean of three independent experiments performed once are shown with error bars representing the SEM (Extended Data Fig. 9).

Purified DMCH-bound mTAAR7f was mixed with mini-G_s_ heterotrimer and Nb35 to form a complex that was isolated from the unbound G protein by size exclusion chromatography (Extended Data Fig. 2a,b) and vitrified for single-particle cryo-EM analysis (Extended Data Fig. 2c,d). The reconstruction of the mTAAR7f-G_s_ complex had a nominal resolution of 2.9 Å (Fig. 1b,c, Extended Data Fig. 2e,f and Extended Data Table 1). The receptor portion of the complex was flexible and therefore focussed refinement was used to improve its resolution from 3.5 Å to 3.2 Å (Extended Data Fig. 2e,f and Extended Data Fig. 3). The density of the ligand was clearly distinguishable (Fig. 1c) and the planar configuration of the ligand was observed by the density’s flattened oval shape. Due to the lack of features to guide the exact position of the substituent group, the ligand was placed according to its known functional properties^12^, with the positively charged tertiary amine group adjacent to the carboxylate of Asp127^3.32^ (superscript refers to the Ballesteros-Weinstein numbering system). This position was corroborated by molecular dynamics (MD) simulations (Fig. 1d).

**Figure 3.**
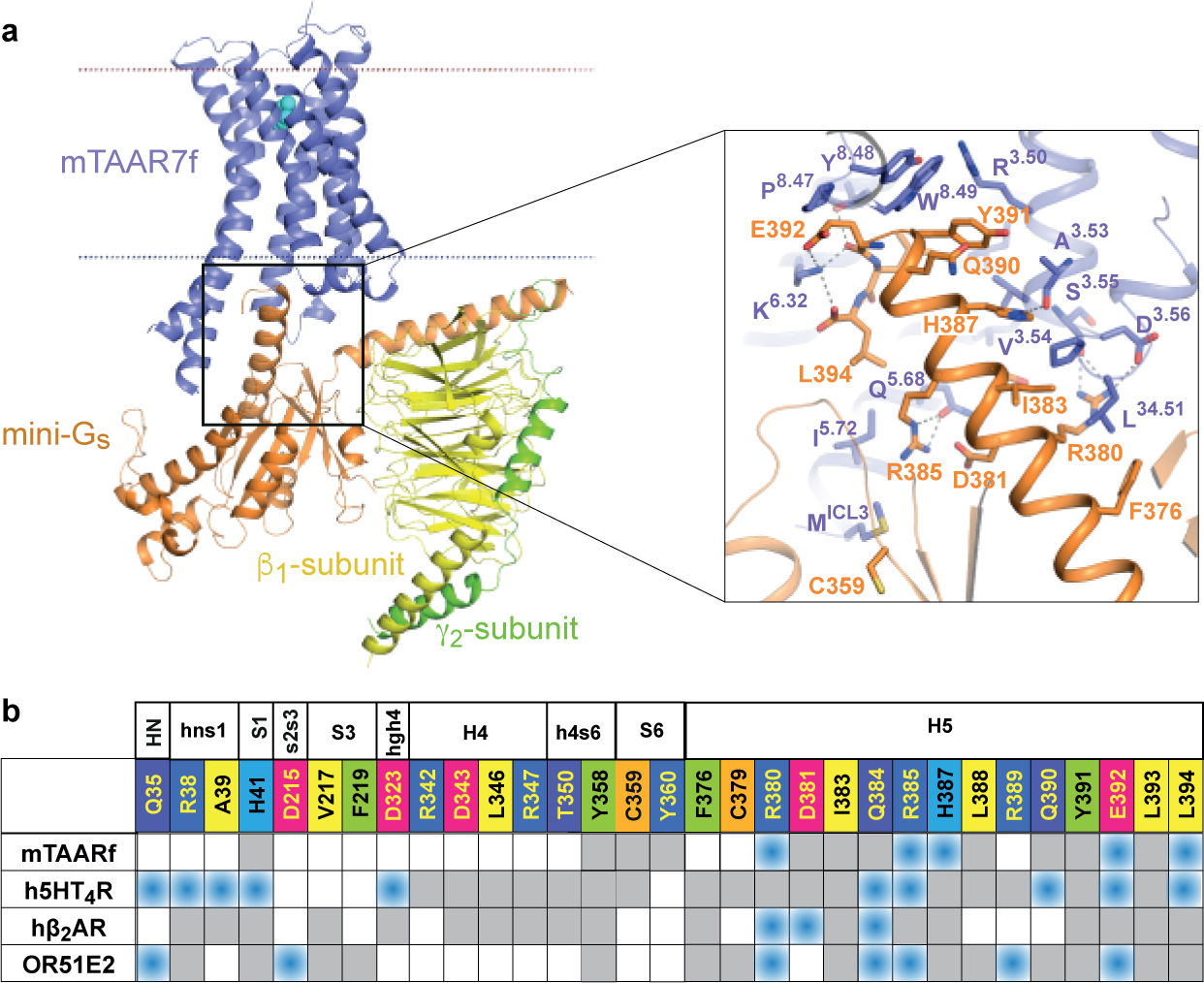
Interactions between mini-G_s_ and mTAAR7f. **a**, Cartoon of the mTAAR7f-G_s_ complex with an inset highlighting interactions between the α5 helix of mini-G_s_ and mTAAR7f (distance cut-off ≤ 3.9 Å). **b,** Comparison of amino acid contacts (distance cut-off ≤ 3.9 Å) made by the α-subunit of G_s_ and mTAAR7f, h5HT_4_R (PDB 7XT8), hβ_2_AR (PDB 3SN6) and OR51E2 (PDB 8F76); blue, polar contacts; grey, van der Waals contacts.

### Overall architecture of mTAAR7f and the orthosteric binding site

The overall structure of mTAAR7f resembles the canonical structure of Class A GPCRs coupled to a G protein. Alignment of mTAAR7f with the most closely related receptors by a phylogenetic analysis of their amino acid sequences showed greatest similarities with β_2_AR (Extended Data Fig. 4) and the serotonin 5-HT_4_ receptor (5-HT_4_R) with RMSDs (all Cα atoms) of 1.9 Å and 2.7 Å, respectively, for receptors in their G protein-coupled state. In contrast, there is little similarity between mTAAR7f and the only other olfactory receptor structure OR51E2^9^ (RMSD of 5.0 Å, all Cα atoms), which is also in an active state coupled to mini-G_s_, as this is more distantly related phylogenetically (Fig. 1a).

**Figure 4.**
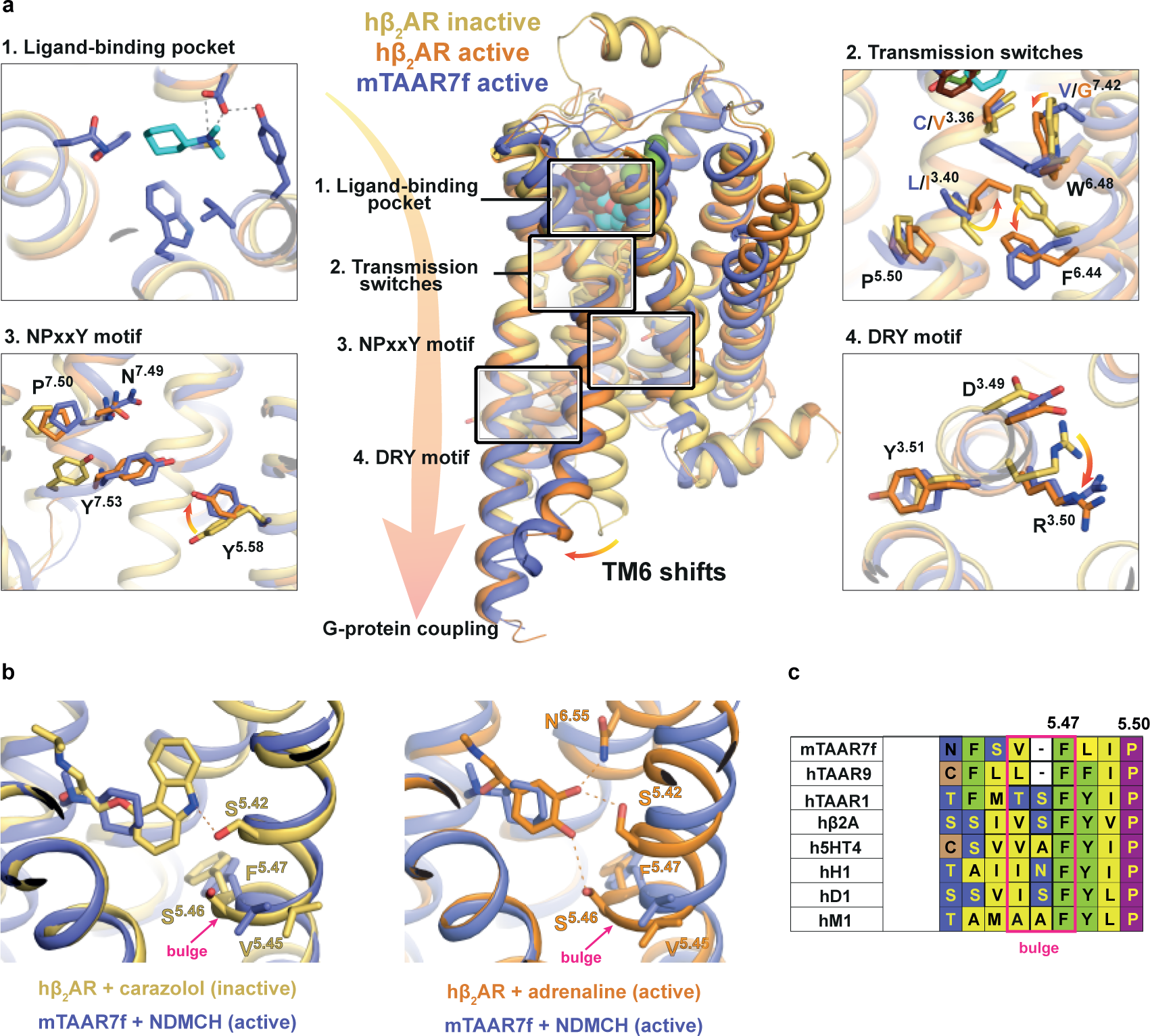
Activation switches in mTAAR7f and β_2_AR. **a**, Conformational changes in the functional motifs are depicted in an alignment of the inactive state of hβ_2_AR (yellow, carazolol-bound, PDB 2RH1), an active state of hβ_2_AR (orange, BI-167107-bound, PDB 3SN6) and mTAAR7 (purple). **b**, Increase in the TM5 bulge in β_2_AR upon the transition from an inactive state (left panel, yellow, PDB 2RH1) to the active state (right panel, orange, PDB 4LDO). Both structures are aligned with the active structure of mTAAR7f-G_s_-DMCH (purple). Hydrogen bonds between the receptors and their corresponding ligands are shown as dashed lines. **c**, Alignment of amino acid residues in the bulge region of aminergic GPCR representatives with mTAAR7f, hTAAR9 and hTAAR1. One amino acid in the bulge region is absent in mTAAR7f and hTAAR9.

The position of the orthosteric binding site (OBS) resembles closely that of the aminergic receptors and not that of OR51E2 (Fig. 2a). In mTAAR7f, the agonist DMCH is found in a cavity formed by transmembrane helices TM3, TM5, TM6 and TM7, and is separated from the outside of the cell by extracellular loop (ECL) 2 (Fig. 2a,d), which is held in position across the OBS by the Class A canonical disulphide bond between Cys205^ECL2^ and Cys120^3.25^. The OBS of mTAAR7f overlaps the positions of the OBS in 5-HT_4_R and β_2_AR and the position of the agonists also overlap (Fig. 2a,c), but the pocket itself is smaller and lacks the extracellular access seen in the aminergic receptors. In contrast, the even smaller binding pocket of propionate in the OR51E2 structure and the position of the agonists do not overlap at all with mTAAR7f, despite sharing the same occluded architecture.

All the receptor-ligand contacts (≤ 3.9 Å; Fig 2d) in mTAAR7f are mediated by eight amino acid residue side chains, four of which are aromatic (Tyr132^3.37^, Trp286^6.48^, Tyr289^6.51^, Tyr316^7.43^), two hydrophobic (Val312^7.39^, Val315^7.42^) and two polar (Asp127^3.32^, Asn217^5.42^). All of the interactions are mediated by van der Waals interactions with the exception of a strong polar interaction between the charges on Asp127^3.32^ and the tertiary amine in DMCH. In MD simulations (Fig. 2e), this interaction was preserved 100% of the time (5 simulations, 1 μs each, 50,000 snap shots per simulation). Other receptor-ligand interactions identified in the cryo-EM structure mediated by van der Waals contacts are present 30-90% of the time (Fig 2e). In addition, the MD simulations identified three other residues that make contact to the ligand 20-70% of the time (Val128^3.33^, Cys131^3.36^, Phe290^6.52^; Fig. 2e). Mutagenesis of residues predicted to make contact to the ligand (either from the cryo-EM structure or from MD simulations) significantly decreases G protein recruitment (Fig. 2f). Of the residues in the OBS of mTAAR7f, Asp127^3.32^, Trp286^6.48^ and Tyr316^7.43^ are all absolutely conserved in all murine and human TAARs (Extended Data Fig. 5b).

Comparison between residues involved in receptor-ligand contacts in mTAAR7f and the aminergic receptors β_2_AR, 5-HT_4_R and 5-HT_1_R highlight commonalities and differences. Three conserved residues (Asp127^3.32^, Trp286^6.48^, Tyr316^7.43^) make contacts to the respective agonists in all four structures (Fig. 2c), with an additional three residues always making contacts (positions 3.37, 5.42, 6.51) and one residue often making contacts (position 7.39). The interaction between DMCH and Asp127^3.32^ is particularly striking as this residue makes interactions with a nitrogen atom in ligands binding to GPCRs throughout the aminergic family. ECL2 often makes contacts to ligands in aminergic receptors (Extended Data Fig. 5a), but does not make contacts to DMCH in mTAAR7f. In the structure of OR51E2, the agonist propionate makes contacts to eight amino residue side chains that form a binding site with strong polar attributes due to the presence of five polar side chains (His104^3×33^, His180^45×52^, Gln181^45×53^, Ser258^6×55^, Arg262^6×59^) and only three hydrophobic side chains (Phe155^4×57^, Leu158^4×60^, Ile202^5×43^)^9^. This is distinct from the predominantly hydrophobic OBS in mTAAR7f. None of the ligand-binding residues in OR51E2 correlate with ligand-binding residues in mTAAR7f, although four of the residues (positions 3×33, 45×52, 5×43 and 6×55) correspond to ligand-binding residues in aminergic receptors, including a residue from ECL2. MD simulations of mTAAR7f indicate that the ECLs are dynamic and allow rapid binding of DMCH (within 100-200 ns, four out of five trajectories, (see Methods; Extended Data Fig. 6a,b), but none of the ECL residues are involved in interacting with the ligand in any of its lowest energy states.

Previous structure-activity relationship (SAR) data for mTAAR7f suggests that ligand binding is highly dependent on the ligand shape and the length of the hydrophobic chain, with aliphatic chains containing less than six carbon atoms being unable to activate the receptor^12^. The size and shape of the OBS seen in the cryo-EM structure clearly imposes restrictions on which ligands can bind. The mTAAR7f mutant Y132C^3.37^ was predicted to expand the size of the OBS and to allow binding of bulkier ligands that activate mTAAR7e which contains a Cys residue at this position; the mutation did indeed reverse the ligand selectivity of the two receptors as predicted^12^ and is consistent with the cryo-EM structure, as a smaller residue at this position would allow ligands to pack between TM3 and TM5.

### G protein coupling interface

The position of the heterotrimeric mini-G_s_ protein in relation to mTAAR7f is similar to that in other class A GPCRs^15^. However, compared to its nearest homologues β_2_AR and 5-HT_4_R, and to OR51E2, mTAAR7f forms fewer contacts between the receptor and the α-subunit of the G protein (Fig. 3a,b). Seventeen residues in the α-subunit make contacts to the receptor, thirteen of which are in the α5 helix and are conserved in other G_s_-coupled receptor structures. The amino acid identity of all the α-subunit residues in contact with the receptor are 100% conserved between G_s_ and G_olf_. The area of the mTAAR7f-G_s_ interface is smaller (1140 Å^2^) compared to that in the 5-HT_4_R-G_s_ (1580 Å^2^) or β_2_AR-G_s_ (1260 Å^2^) complexes.

### Ligand-induced activation of mTAAR7f

The activation of class A GPCRs by diffusible ligands occurs through a series of structural changes commencing with agonist binding, followed often by a contraction of the OBS and then propagation of structural changes through the receptor to the intracellular G protein binding interface (Fig. 4a,b). The resulting outward shift of the intracellular end of TM6 enables coupling of a G protein, as exemplified by the active state of the β_2_AR^16^. The orientation of mTAAR7f transmembrane helices in the cryo-EM structure aligns well with the G protein-coupled active state of β_2_AR (Fig. 4a,b). In addition, mTAAR7f contains hallmarks of activation in the conserved regions essential for stabilisation of the active state, including the P^5.50^-I^3.40^-F^6.44^ motif, the C^3.36^-W^6.48^-x-F^6.44^ motif, the D^3.49^-R^3.50^-Y^3.51^ motif, and the N^7.49^P^7.50^xxY^7.53^ motif (Fig 4a, Extended Data Fig. 5c). The ionic lock between Arg^3.50^ and Asp^3.49^ is a hallmark of an inactive state of Class A receptors, which is broken upon receptor activation through a rotamer change of Arg^3.50^. In the mTAAR7f structure, the positions of Arg145^3.50^ and Asp144^3.49^ are identical to the equivalent residues in the active state of β_2_AR (Fig. 4a). Similarly, the positions in mTAAR7f of Tyr326^7.53^ in the NPxxY motif and the associated Tyr232^5.58^ align well with the equivalent residues in the active state of β_2_AR and not the inactive state. However, only portions of the CWF and PIF motifs follow the canonical pattern of rotamer conformations observed in β_2_AR (Fig. 4a). Phe282^6.44^ in the PIF motif in mTAAR7f does align well with the respective rotamer in β_2_AR, but Leu135^3.40^ in mTAAR7f cannot adopt the active conformation of Ile^3.40^ in β_2_AR due to the position of Trp286^6.48^. The position of Trp286^6.48^ in mTAAR7f is rotated by 35° around the TM6 helical axis compared to its position in β_2_AR, resulting in a 4.2 Å difference in its position (measured at the CH2 atom). The shift of Trp286^6.48^ in mTAAR7f also causes Phe^6.44^ to adopt an active state conformation to prevent a clash. The position of the highly conserved Trp^6.48^ in Class A GPCRs has been described as one of the key elements of activation of many GPCRs^17^, making this a likely candidate in the activation of mTAAR7f.

Why does Trp286^6.48^ adopt such an extreme conformation compared to β_2_AR? The ligand DMCH makes van der Waals contacts to Trp286^6.48^ and this could be one reason why it is shifted greatly compared to its position in β_2_AR. The rotations of DMCH observed in the MD simulations would place Trp286^6.48^ in this position and this is evident from the Chi2 dihedral angle fluctuations being far lower when DMCH is bound compared to when it is not (compare Step 2 with Step 3 in the ligand binding pathway, Extended Data Fig. 6a). The rotamer of Trp286^6.48^ is directly impacted by the DMCH and is the last step in the ligand binding process observed by MD (Step 4, Extended Data Fig. 6a). Another residue that may play role in the position of Trp286^6.48^ is Val315^7.42^ which is only 4 Å from Trp^6.48^ in mTAAR7f, and would clash if Trp286^6.48^ were to adopt the active state conformation observed in β_2_AR. The importance of these two residues in the activation of mTAAR7f is apparent from mutagenesis data. The mutants W268Y^6.48^ and V315A^7.42^ both show significantly decreased E_max_ and EC_50_ for activation compared to the wild type receptor (Fig. 2f).

Comparisons between mTAAR7f and β_2_AR can also help to formulate a mechanism of how DMCH binding may potentially activate the receptor. In β_1_AR and β_2_AR, ligand-induced activation is caused by the para-hydroxyl of the catecholamine moiety of the agonist inducing a rotamer change of Ser^5.46^ and the contraction of the OBS by 1-2 Å^18,19^. Coupling of the G protein causes a further contraction of the OBS, predominantly through the movement of the extracellular ends of TM6 and TM7, resulting in decreased on/off rates of the ligand and an increase in agonist affinity due to an increased number and/or strength of ligand-receptor interactions^20,21^. mTAAR7f differs from the βARs in that there is only a weak van der Waals interaction between the agonist and TM5 (Asn217^5.42^), and also that the characteristic bulge formation upon ligand activation of βARs is absent. Amino acid sequence alignments between aminergic receptors and TAARs show that there is a one amino acid deletion in this region in the TAARs (Fig 4c), leading to TM5 being unable to form a bulge. Therefore, it is likely that the activation cascade upon ligand binding to mTAAR7f differs subtly to that of the βARs.

Based on the active-state mTAAR7f structure and our extensive knowledge of the activation of the βARs, we suggest here a possible mechanism of ligand activation of m7TAARf. Binding of DMCH occurs predominantly through charge-charge interactions between the tertiary amine of the ligand and Asp127^3.32^, and extensive van der Waals interactions with the hydrophobic OBS. This causes a contraction of the OBS through interactions between DMCH and residues in TM6 and TM7 (Val312^7.39^, Tyr316^7.43^, Val315^7.42^, Y289^6.51^, Trp286^6.48^) and the stabilisation of the interaction between TM3 and TM7 by a hydrogen bond between Tyr316^7.43^ and Asp127^3.32^. The position of the DMCH and Val315^7.42^ causes Trp286^6.48^ to rotate and induce activation of downstream activation motifs (PIF, NPxxY and DRY), ultimately resulting in the outward movement of TM6 and G protein coupling. Of course, in the absence of an inactive state structure of mTAAR7f this is currently a working hypothesis, but it is supported by both mutagenesis data and MD simulations. The mutants W286Y^6.48^ and Val315^7.42^ both show low levels of DMCH-induced G protein coupling, consistent with their roles in receptor activation. The mutant D127A^3.32^ significantly decreases agonist-induced G protein coupling, and the mutant C131A^3.36^ has a similar effect; the structure suggests that C131A^3.36^ is important in maintaining the rotamer of Asp127^3.32^ for optimal binding to DMCH. Other mutations (Y132A^3.37^, Y289A^6.51^) also reduce agonist-induced signalling, probably by reducing the strength of DMCH-mTAAR7f interactions.

Full atomistic MD simulations are inadequate to observe the full transition between an inactive state of a GPCR to an active state. In the simulations performed here to look at the movements of residues and secondary structure within mTAAR7f in the absence of the G protein and/or DMCH, we analysed overall trends in the context of deactivation. Five simulations were performed (1 μsec each) either on TAAR7f-DMCH-mini-G_s_, TAAR7f-DMCH or TAAR7f (see methods and Extended Data Fig. 7a-c). In the absence of G protein, the mean GPCR backbone RMSD increased as expected due to the lack of strong stabilisation of the GPCR active conformation through G protein allosteric coupling (Extended Data Fig. 7a)^20,21^. In addition, the simulations show that the OBS increases in volume when ligand and G protein are removed, which is consistent with receptor deactivation. Observation of the activation microswitches in the simulations (Extended Data Fig. 7b,c) also indicated that they all started to move towards inactive state conformations, as assessed by measuring distances between specific pairs of residues; similar results were observed for β_2_AR (Extended Data Figure 7c). In contrast, an analogous analysis on OR51E2 showed a distinct series of changes upon removal of the agonist and G protein^9^, suggesting that the deactivation process is different from mTAAR7f and β_2_AR.

### Addendum

During the preparation of this manuscript structures of mouse TAAR9 (mTAAR9) were published bound to three different ligands, including DMCH^22^. The amino acid sequences of mTAAR9 and mTAAR7f are similar (69% identical) and the structures are also similar (overall RMSD 1.2 Å). The pose of the DMCH in the OBS of mTAAR9 is different from what we observe with the cyclohexylamine ring rotated by 55° around an axis defined between the tertiary amine and cyclohexylamine ring; this pose probably resulted from the significantly worse density for the ligand in the hTAAR9 structure, but the position is in line with the range of poses we observe during MD simulations. The authors’ analysis of the activation mechanism based on mutational analysis broadly agrees with ours presented here, namely that the mechanism is most similar to that of β_2_AR and that Trp^6.48^ plays an important role in ligand-induced G protein coupling.

## Acknowledgements

We thank R. Axel, S. Liberles, V. Velazhahan, K. Yamashita, G. Murshudov, B.Ahsan, and T. Warne for helpful discussions. We acknowledge the MRC Laboratory of Molecular Biology Electron Microscopy Facility for access and support of electron microscopy sample preparation and data collection, the LMB scientific computing for technical support and the Flow Cytometry facility for support in cell analysis. We also thank S. Liberles for supplying original clones of receptors. The work in C.G.T.’s laboratory was supported by core funding from the Medical Research Council [MRC U105197215] and by a grant from Sosei Heptares. Work in the lab of F.M. was funded by the National Institutes for Health (USA) grant GM132120. S.N.W.’s lab was funded by a Sir Henry Dale fellowship from the Wellcome Trust and the Royal Society London (Grant Number 101234/Z/13/Z) and by the Isaac Newton Trust Cambridge (Grant Number 15.40(a)). The work in N.V.’s lab was funded by grants from the National Institutes of Health (2R01-GM117923). A.N.K. is supported by the BBSRC Doctoral Training Programme at the University of Nottingham.

## Author contributions

A.G. optimized the constructs, performed receptor expression, purification, preparation of cryo-EM grids, cryo-EM data collection, data processing, structure determination and model building. F.H. developed the initial expression and purification strategy. P.C.E. performed the expression and purification of G protein. Q.C. cloned receptor variants for functional studies. Y.L. advised on receptor expression, purification and data collection, supervised data processing, structure determination and performed model building. E.M. and N.M. performed the molecular dynamics simulations, and N.M. and N.V. did the analysis of the MD simulation trajectories. A.N.K. and E.K. performed G protein-coupling assays, and A.N.K., E.K. and D.V. did the analysis of the data. J.K. and F.M. contributed significantly at the initial stages of receptor screening and selection. A.G. and C.G.T carried out structure analysis and manuscript preparation. C.G.T. and S.N.W. managed the overall project. The manuscript was written by A.G. and C.G.T., and included contributions from all the authors.

## Author information

Reprints and permissions information is available at www.nature.com/reprints. The authors declare the following competing interests: CGT is a shareholder, consultant and member of the Scientific Advisory Board of Sosei Heptares. Correspondence and requests for materials should be addressed to.

## Data availability statement

Structures have been deposited in the Protein Data Bank (PDB; https://www.rcsb.org/), and the associated cryo-EM data has been deposited in the Electron Microscopy Data Bank (EMDB; https://www.ebi.ac.uk/pdbe/emdb/) and the Electron Microscopy Public Image Archive (EMPIAR; https://www.ebi.ac.uk/empiar/): PDB XXXX, EMDB-XXXXX, EMPIAR-XXXXX. There are no restrictions on data availability.

## Materials and Methods

### Expression and purification of the mini-G_s_ heterotrimer and Nb35

The components of the heterotrimeric G protein (mini-G_s_ construct 399, β_1_-subunit, γ_2_-subunit and Nb35) were expressed and purified as described previously^18,24,25^. In brief, mini-G_s_ in plasmid pET15b was expressed in bacterial strain BL21-CodonPlus(DE3)-RIL. His-tagged protein was purified via Ni^2+^-affinity chromatography, followed by cleavage of the histidine tag using TEV protease and negative purification on Ni^2+^-NTA to remove the TEV and undigested mini-G_s_. β_1_ and unlipidated (C68S mutation) γ_2_ subunits were co-expressed in HighFive (*Trichoplusia ni*) cells (Expression Systems; we did not test for mycoplasma). The protein was purified via Ni^2+^-affinity chromatography followed by anion exchange chromatography. Aggregates in the purified β_1_γ_2_ complex were removed by size-exclusion chromatography (SEC). The three G protein subunits were mixed and the heterotrimeric G protein isolated by SEC, concentrated, aliquoted and flash-frozen in liquid nitrogen until further use. Nanobody-35 (Nb35) was expressed from plasmid pET26b in the periplasm of *E. coli* strain BL21-CodonPlus(DE3)-RIL, extracted, and purified by Ni^2+^-affinity chromatography, according to previously described methods, followed by ion exchange chromatography^18^. Purified Nb35 was concentrated and flash frozen in liquid nitrogen until further use.

### Cloning and expression of the mTAAR7f

The wild type murine TAAR7f gene (UniProt Q5QD08) was synthesized and cloned into a modified pFastBac1 vector with HA signal sequence, FLAG tag, 10x His-tag and TEV protease cleavage site before the receptor N-terminus and HRV 3C cleavage site followed by eGFP after its C-terminus. Cloning was performed by overlap extension PCR using *Escherichia coli* DH10B cells (Thermo Fischer) and positive clones identified by DNA sequencing. High titre (>3 × 10^8^ viral particles per ml) recombinant baculovirus was obtained using the Bac-to-Bac expression system (Invitrogen) in Sf9 (*Spodoptera frugiperda*) cells grown in Sf-900 II medium (Thermo Fischer) and its titre was checked with flow cytometry technique using anti-gp64 conjugated antibodies^26^. *Trichoplusia ni* High Five cells (Thermo Fisher Scientific; we did not test for mycoplasma) were grown in suspension in ESF921 media (Expression Systems) and infected at a density of 2-3 million cells per ml using a multiplicity of infection of 7-10. The cells were then collected by centrifugation and resuspended in m7-glycerol+ buffer (20 mM HEPES/KOH pH7.5, 150 mM potassium chloride, 10 mM sodium chloride, 10 mM magnesium chloride, 20% v/v glycerol) supplemented with 1 tablet / 50 ml Complete protease inhibitor (Roche), 1 mM PMSF), flash-frozen in liquid nitrogen, and stored for several months at −80 °C until further use.

### Purification of mTAAR7f

Cells were thawed and lysed by two washes in low salt buffer (25 mM Na HEPES pH 7.5, 1 mM EDTA, Complete protease inhibitor, 1 mM PMSF) followed by two washes in high salt buffer (25 mM Na HEPES pH 7.5, 1 mM EDTA, 1M NaCl, Complete protease inhibitor, 1 mM PMSF), followed by one wash in the low salt buffer. During each round, the pellets were resuspended using an UltraTurrax homogeniser and centrifuged (235000 x*g*, 60 min 4°C). The pellets were resuspended in m7-glycerol+ buffer supplemented with PMSF and Complete protease inhibitor and flash frozen in liquid nitrogen.

Previously frozen cell membranes containing overexpressed mTAAR7f receptor were thawed and resuspended to a final volume of 160 ml in m7-glycerol+ buffer which was supplemented with 2 mg/ml of iodoacetamide, 6.7 mM N,N-dimethylcyclohexylamine (DMCH) and EDTA-free complete protease inhibitor tablets (Roche). The mixture was incubated at 4°C with rotation for 2 hours. LMNG/CHS mixture (5/0.5 % w/v stock) was added to the final concentration of 1/0.1% LMNG/CHS to solubilise the receptor. (4 °C, 1 hour) and then centrifuged (430,000 x*g*, 1.5 hours, 4°C). The supernatant was incubated overnight with a 2 ml bed volume of Super Ni-NTA Affinity Resin (ProteinArk) and 20 mM imidazole at 4°C with rotation. All further purification steps were performed at 4 °C. The following day the resin was placed into an empty PD10 gravity column and washed with 10 ml of buffer m7 supplemented with 8 mM ATP, 20 mM imidazole, 6.7 mM DMCH and 0,01/0.001% LMNG/CHS. The resin was further washed with 15 ml of buffer m7 supplemented with 40 mM imidazole, 6.7 mM DMCH and 0.01/0.001% LMNG/CHS. The receptor was eluted with elution buffer containing 20 mM HEPES/KOH pH7.5, 150 mM potassium chloride, 10 mM sodium chloride, 10 mM magnesium chloride, 20% v/v glycerol, 300 mM imidazole, 6.7 mM DMCH and 0,01/0.001% LMNG/CHS. Eluted fractions were pooled and concentrated in a 50 kDa molecular weight cut-off Amicon ultracentrifugal concentrator (Merck) at 2,000 x*g* and exchanged into the same buffer used for elution (without imidazole) using a PD10 desalting column. To cleave off the His tag, the protein was incubated overnight with TEV protease in the presence of 0.5 mM DTT.

The purity of mTAAR7f was then improved significantly using a reverse Ni-NTA purification step of the TEV cleaved mixture by incubating it with rotation for 2 hours with 0.5 ml bed volume of Super Ni-NTA Affinity Resin (ProteinArk) supplemented with 10 mM imidazole. eGFP was then cleaved off by incubation with HRV 3C protease in the presence of 0.5 mM DTT. The cleaved receptor was separated from eGFP by SEC on a Superdex 200 10/300 GL column (GE Healthcare) pre-equilibrated with m7-glycerol+ buffer containing 6.7 mM DMCH. Peak fractions were pooled and concentrated using a 50 kDa molecular weight cut-off Amicon ultracentrifugal concentrator (Merck).

### mTAAR7f - miniGs399 - β_1_ - γ_2_ - Nb35 complex assembly

Purified and concentrated mTAAR7f was mixed with about 10x molar excess of both the heterotrimeric G protein and Nb35 and 0.5 U apyrase (NEB) and incubated overnight. The following morning unbound G protein and Nb35 were separated from the complex by SEC on a Superdex 200 10/300 GL column (GE Healthcare) pre-equilibrated with the buffer containing 20 mM HEPES/KOH pH7.5, 150 mM KCl, 10 mM NaCl, 10 mM MgCl_2_, 6.7 mM DMCH. Fractions corresponding to the size of the complex were pooled, and concentrated in a 100 kDa cut-off concentrator.

### Grid preparation of the complex and data collection

Grids for cryo-EM (UltrAuFoil 1.2/1.3) were prepared by applying 3 μl sample concentrated to 0.9 mg/ml on a glow-discharged grid (2 min in Ar-Oxy 9-1 plasma chamber, at Forward Power of 38 W, Reflected Power of 2W; Fischione). The excess sample was removed by blotting for 3 s before plunge-freezing in liquid ethane (cooled to −181 °C) using a FEI Vitrobot Mark IV maintained at 100% relative humidity and 4 °C. Data was collected in-house from a single grid on the FEI Titan Krios microscope at 300 kV equipped with a Falcon 4 detector in counting mode. A total of 12273 movies were collected in one session with a fluence of 55 *e*^−^/Å^2^ at 96,000x magnification (0.824 Å/pixel). The gain reference file was provided by the facility and used unmodified.

### Cryo-EM data processing

12273 movies in .EER format were converted into .tiff format with relion_convert_to_tiff utility^27,28^, grouping the frames to get the dose per frame of 1.38 e^−^/Å^2^. The resulting movie stack was imported into CryoSparc v4.1.1+patch230110^29^ and the processing was performed there unless specified otherwise (Extended Data Fig. 3). Overall, drift, beam-induced motion and dose weighting were corrected with Patch Motion Corr. CTF fitting estimation were performed using Patch CTF estimation. The exposures were manually curated: the only images kept had an estimated CTF resolution of <5 Å, motion distance <200 pixels and no obvious outliers in terms of estimated relative ice thickness. This yielded a stack of 11,157 movies. Auto-picking was performed with Gaussian circular and elliptical blobs as templates with inner and outer diameters 80 and 160 Å respectively and 0.5 diameters as a minimum separation distance. Particle picks were curated to remove obvious junk peaks (e.g. the ones outside of foil holes or on contaminants) and then extracted with the box size of 300 Å and down-sampled to 1.648 Å/pixel. The particles were subjected to five rounds of 2D classification and the clean particle stack was re-extracted at 0.824 Å/pixel.

*Ab initio* reconstruction for the mTAAR7f-G protein complex was made using 198,414 particles belonging to clean 2D classes with different orientations. Hetero refinement was performed with the *ab initio* reconstruction of the receptor complex and three noise classes as input. The good class from hetero refinement was subjected to one round of non-uniform refinement resulting in an initial 3D volume.

The curated particle image coordinate data was exported from cryoSPARC using pyem v0.5^30^. Beam-induced motion correction and dose-weighting were repeated using RELION’s implementation of motion correction with a 5×5 patch array. Particle images were then re-extracted from the averaged micrographs and realigned to the consensus map through non-uniform refinement. Coordinates and transformations were exported with pyem for Bayesian polishing in RELION, maintaining image dimensions of the shiny particle stack.

Per-hole beam image-shift exposure groups were identified with EPU_group_AFIS^31^. Particle images were assigned to exposure groups and refined in cryoSPARC using non-uniform refinement^32^, iterated with particle defocus refinement and higher order CTF refinement (beam tilt and trefoil parameters)^33^, to an estimated global resolution of 2.92 Å (gold-standard FSC=0.143; Extended Data Figs. 2e,f).

To help receptor modelling, especially its most flexible regions (helices 1 and 2) focused refinement was performed using a mask on the receptor, which visually improved the map quality. This refinement centred on a mask of the receptor region, using pose/shift Gaussian priors and 3°/3Å standard deviations, yielded a 3.2Å focused map. Local resolution estimation was performed using cryoSPARC’s adaptive window implementation. The nominal local resolution for the receptor was improved from 3.5Å in the overall consensus map to 3.2Å in the focused map. Local sharpening was performed with LocalDeblur^34^ using half-maps and estimated local resolution maps.

### Model building and refinement

Initial models of heterotrimeric mini-G_s_ and Nb35 were sourced from PDB 7T9I. A *de novo* model of TAAR7f was generated from the focused map and protein sequence using ModelAngelo^35^. Overall, this initial model agreed with the map except for poorly resolved regions which were added and iteratively modelled afterwards. Chemical restraints for N,N-dimethylcyclohexylamine were generated using phenix.eLBOW (AM1 optimisation)^36^ and manually fitted into the density. Manual rebuilding was performed in COOT^37^ and ISOLDE^38^ (in ChimeraX^39^) and further refined against the locally-sharpened consensus map using phenix.real_space_refine. 3D variability analysis^40^ was performed on the consensus map using the refinement mask and at a filter resolution of 4 Å with a high-pass prior of 20 Å.

### Molecular dynamics simulations

Three distinct molecular dynamics simulations were performed to investigate TAAR7f behaviour under different conditions: (1) an apo state simulation, excluding both ligand and G protein; (2) a ligand-bound simulation, incorporating the ligand but devoid of the G protein; (3) a G protein-bound simulation, including the ligand and mini-G_s_, but omitting the β-subunit and γ-subunit of the heterotrimeric G protein and Nb35. The starting point for these simulations was the cryo-EM structure of the DMCH-bound TAAR7f-G_s_ complex. TAAR7f in each simulation was encapsulated within a layer of cholesteryl hemisuccinate, employing the same methodology as that used in our prior OR study^9^. Utilizing the membrane builder module^41^ from CHARMM-GUI^42^, the simulation system was assembled by embedding the complexes into a POPC bilayer with X-Y dimensions of 85 Å - 85 Å for apo and ligand-bound systems, and 125 Å - 125 Å for the G protein-bound system. The resulting system was then immersed into a TIP3 water box providing a 10 Å margin along the Z-axis from the protein surface (Z dimension ∼ 125 Å), followed by neutralization using 0.15 M NaCl. The simulations systems were parameterized by CHARMM36m force field^43^, and all simulations were performed by GROMCS-2022 version^44^.

An additional MD simulation was established to scrutinize the ligand association process. This involved positioning apo state TAAR7f (no G protein involved) within a grid box with 8 * 8 * 8 units of DMCH, with a grid spacing (distance between any two adjacent DMCH residues) of 12 Å. This complex was configured into a simulation box by adopting the same procedure previously outlined, after the removal of any DMCH molecules found to overlap with the POPC bilayer.

The simulation systems were progressively heated from 0 K to 310 K using a constant volume-constant temperature (NVT) ensemble and a Nosé-Hoover thermostat^45^. This was followed by a 30 ns equilibration protocol implemented using a constant pressure-constant temperature (NPT) ensemble. Throughout both heating and equilibration phases, harmonic positional restraints were applied to proteins, the ligand, and the heavy atoms of the head group of the cholesteryl hemisuccinate and POPC lipids. The system was initiated with a positional restraint force constant of 10 kcal/mol-Å^2^, which was upheld for a duration of 5 ns. This was succeeded by a decrease in the constant to 5 kcal/mol-Å^2^, which was maintained throughout the next 5 ns. The constant was then methodically reduced to 0 kcal/mol-Å^2^ at a decrement rate of 1 kcal/mol-Å^2^ for each successive 5 ns span. The concluding phase of the equilibration process was carried out with a restraint constant of 0 kcal/mol-Å^2^ over a 10 ns period. Pressure control was facilitated by the Parrinello-Rahman method^46^, and the simulation system was harmonized with a 1 bar pressure bath. The concluding snapshot from the equilibration stage was chosen as the commencement conformation for five unrestrained NPT simulation runs, each with distinct random seeds. Each of these runs spanned 1000 ns at a temperature of 310 K. In the case of the ligand association simulations, the duration of each run was extended to 2200 ns. In all the simulations, the LINCS algorithm was utilized for all water bonds and angles, with a time step of 2 fs for integration. Non-bond interactions were subjected to a cut-off of 12 Å, and the particle mesh Ewald method was used to handle long-range Lennard-Jones interactions^47^. Molecular dynamics snapshots were saved at intervals of every 20 ps.

To ascertain the flexibility of the ligands, we conducted ligand clustering analysis utilizing the cluster-analysis-using-VMD-TCL script (https://github.com/anjibabuIITK/CLUSTER-ANALYSIS-USING-VMD-TCL). The merged production trajectories from both the ligand-bound and G protein-bound simulations were initially aligned using the backbone atoms of TAAR7f. Following this, a cut-off value of 1.5 Å was utilized to group ligand conformations based on the root-mean-square deviation (RMSD) of their heavy atoms (Fig. 1d).

To scrutinize the TAAR7f residues establishing stable contacts with the DMCH ligand, we implemented contact frequency analysis utilizing the ‘get_contact’ script (https://getcontacts.github.io/). This analysis was performed on the combined production runs from both the ligand-bound and G protein-bound simulations. TAAR7f and the DMCH ligand were designated as selection 1 and selection 2 respectively. All categories of contacts were considered in the analysis, and the default parameters were employed to evaluate the formation of contacts (Fig. 2e).

Using the MDAnalysis module^48^, we calculated the average ligand RMSD for each of the five individual production runs in both the ligand-bound and G protein-bound simulations. The calculation was performed on the ligand’s heavy atoms, subsequent to aligning the backbone atoms of TAAR7f. This resulted in five average RMSD values for each simulation. We compared these five values, presenting their average and standard deviation in a bar graph (Fig. 1 d). The graph also includes p-values derived from a t-test, offering a statistical comparison between the two simulations.

The calculation of GPCR RMSD was conducted in a similar manner to the ligand RMSD, but utilized the backbone atoms of TAAR7f. No G protein was involved in this calculation. The average TAAR7f RMSD was determined for each individual production run for the Apo, ligand-bound, and G protein-bound simulations, and the results were plotted in a bar graph with accompanying p-values, as depicted in (Extended Data Fig. 7a).

The calculation of ligand binding site volume was conducted using the Maestro SiteMap module (Schrödinger Release 2023-2: SiteMap, Schrödinger, LLC, New York, NY, 2021.). For each production run across all simulations, frames were extracted at the end of both 500 ns and 1000 ns of simulations, resulting in 10 frames per simulation. In the case of the ligand-bound and G protein-bound simulations, the binding pocket was defined by centring on the ligand, and a 6 Å cut-off was applied to establish the binding region. We employed the “standard grid” with a “more restrictive definition of hydrophobicity”, and the site was truncated 4 Å from the nearest site point. In the Apo simulation, the ligand was first docked into the vacant binding site for the 10 frames, and then the same procedure was followed to calculate binding site volume. The ten volume values from each simulation were plotted as a bar graph, with associated p-values (Extended Data Fig. 7a).

The Chi2 dihedral angle of W286 was determined using the MDAnalysis module with the dihedral angle sequence: CA-CB-CG-CD2, designated as the Chi2 angle. Subsequently, these dihedral angles were represented in a histogram constructed using the numpy.hist function, with a bin width of 50 (Extended Data Fig. 7c).

Using MDAnalysis, we determined microswitch distances by calculating the shortest separations between specified atom pairs in various configurations. These include the D127-Y316 distance (between OD1/OD2 atoms of D127 and the OH atom of Y316), the sodium binding site distance (OD1/OD2 atoms of D93 and the OG atom of S134), the NPxxY motif distance (OD1/ND2 atoms of N322 and the OH atom of Y326), and the YY motif distance (OH atoms of Y326 and Y232). This analysis was performed on residues sharing the same Ballesteros-Weinstein numbering in the β_2_AR simulation retrieved from GPCRmd.org (apo state, ID 116; active state, ID 117).

We executed ligand clustering analysis on the four ligand association trajectories using the TICC module from the get_contact script. In these trajectories, different DMCH residues associate with TAAR7f. We limited the association trajectory to include only TAAR7f and the specific DMCH residue in association with TAAR7f, excluding all other non-bound DMCHs. Initially, we computed the contact frequency between the associated DMCH and TAAR7f following the procedure described earlier. We then clustered the frames using the TICC module, arbitrarily setting the cluster count to five. Upon comparison of the representative conformations from each cluster across the four association trajectories, we identified three stable states, which are further elaborated in the results section. Regarding distance measurements, the ligand-D296/D296 distance was quantified as the minimum distance between the N atom of DMCH and the OD1/OD2 atoms of aspartate. On the other hand, the ligand-Y308/W286 distance was determined by measuring the centre of mass distance between the cyclohexane ring of DMCH and the heavy atoms of Y308/W86 (Extended Data Fig. 6).

### Signalling assays

Full length mTAAR7f was cloned into plasmid pcDNA4/TO, encoding a signal peptide, a twin-strep tag, a SNAP-tag at the N-terminus of mTAAR7Ff and nanoluc at the C-terminus. Cloning success was confirmed by Sanger sequencing. NES-Venus-miniG_s_ plasmid^49^ as a kind gift of Nevin Lambert’s lab.

HEK293T/17 cells (ATCC CRL-11268) were transiently transfected with pcDNA4/TO plasmid constructs expressing NES-Venus-mGs and mTAAR7f derivatives. HEK293T/17 cells were cultured in T75 flasks in DMEM, supplemented with 10% FCS, at 37°C in a humidified incubator with 5% CO_2_. After reaching 80%-90% confluency, cells were detached by trypsinization, collected by centrifugation at (350 x*g*, 5 min, 21°C) and resuspended in a fresh media. For PEI (polyethylenimine) transfection, 3 µg of PEI was solubilized in 100 µL OptiMEM (Thermo Fisher Scientific) and was mixed with 1 µg total DNA (100 ng receptor plasmid + 200 ng mGs plasmid + 700 ng salmon sperm DNA (Sigma) solubilized in 100 µL OptiMEM. A cell suspension at 250,000 cells/mL density was prepared and 1 mL of it was added to 200 µL of DNA-PEI mixture. Then, 100 µL aliquots of the transfected cell suspension were seeded in 12 wells of a 96 well white plate with clear flat bottom (25,000 cells per well). After a minimum of 48 hours incubation of plates at 37°C, in a humidified atmosphere with 5% CO_2_, they were checked for at least 60-70% confluency under a microscope. After reaching suitable confluency, the growth media was aspirated, and cells were washed once with 90 µL of assay buffer (HBSS + 0.5% BSA + 0.5mM HEPES pH 7.4 + 0.01% ascorbic acid), which was warmed to 37°C in a water bath. The nanoluc substrate furimazine was added to assay buffer (8 µM final concentration), and 90 µL of this nanoluc substrate containing assay buffer was added to each well. To improve signal level, a white sticker was attached to the bottom and the plates were read in a PHERAstar FSX Microplate reader using an optic module Lum 550-LP 450-80 for 10 min before adding any compound. Finally, 10 µL of compound dilutions were added to the wells and the plate reading was continued for a further 30 min. The data were analysed using GraphPad Prism 9 using standard concentration-response models defined in the software.

## Extended Data Figure legends

**Extended Data Table 1.** Cryo-EM data collection and refinement statistics.

|  |  |
| --- | --- |
| **Data collection and processing** |  |
| Magnification | 96,000x |
| Voltage (kV) | 300 |
| Electron exposure (e<sup>-</sup>/Å<sup>2</sup>) | 55 |
| Defocus range (μm) | -0.8 to -2.4 |
| Pixel size (Å) | 0.824 |
| Symmetry imposed | C1 |
| Initial particle images (no.) | 478,036 |
| Final particle images (no.) | 172,639 |
| Map resolution (Å) | 2.9 |
| FSC threshold | 0.143 |
| Map resolution range (Å) | 2.6 to 5 |
| **Refinement** |  |
| Initial model used (PDB code) | 7T9I |
| Model composition in the ASU |  |
| Non-hydrogen atoms | 7824 |
| Protein residues | 993 |
| Ligands | 37 |
| *B* factors (Å<sup>2</sup>) |  |
| Protein | 94.9 |
| Ligand | 119.9 |
| R.m.s. deviations |  |
| Bond lengths (Å) | 0.004 |
| Bond angles (°) | 0.766 |
| Validation |  |
| MolProbity score | 1.35 |
| Clash score | 3.72 |
| Poor rotamers (%) | 0 |
| Ramachandran plot |  |
| Favoured (%) | 97.58 |
| Allowed (%) | 2.42 |
| Disallowed (%) | 0 |

**Extended Data Fig. 1.**
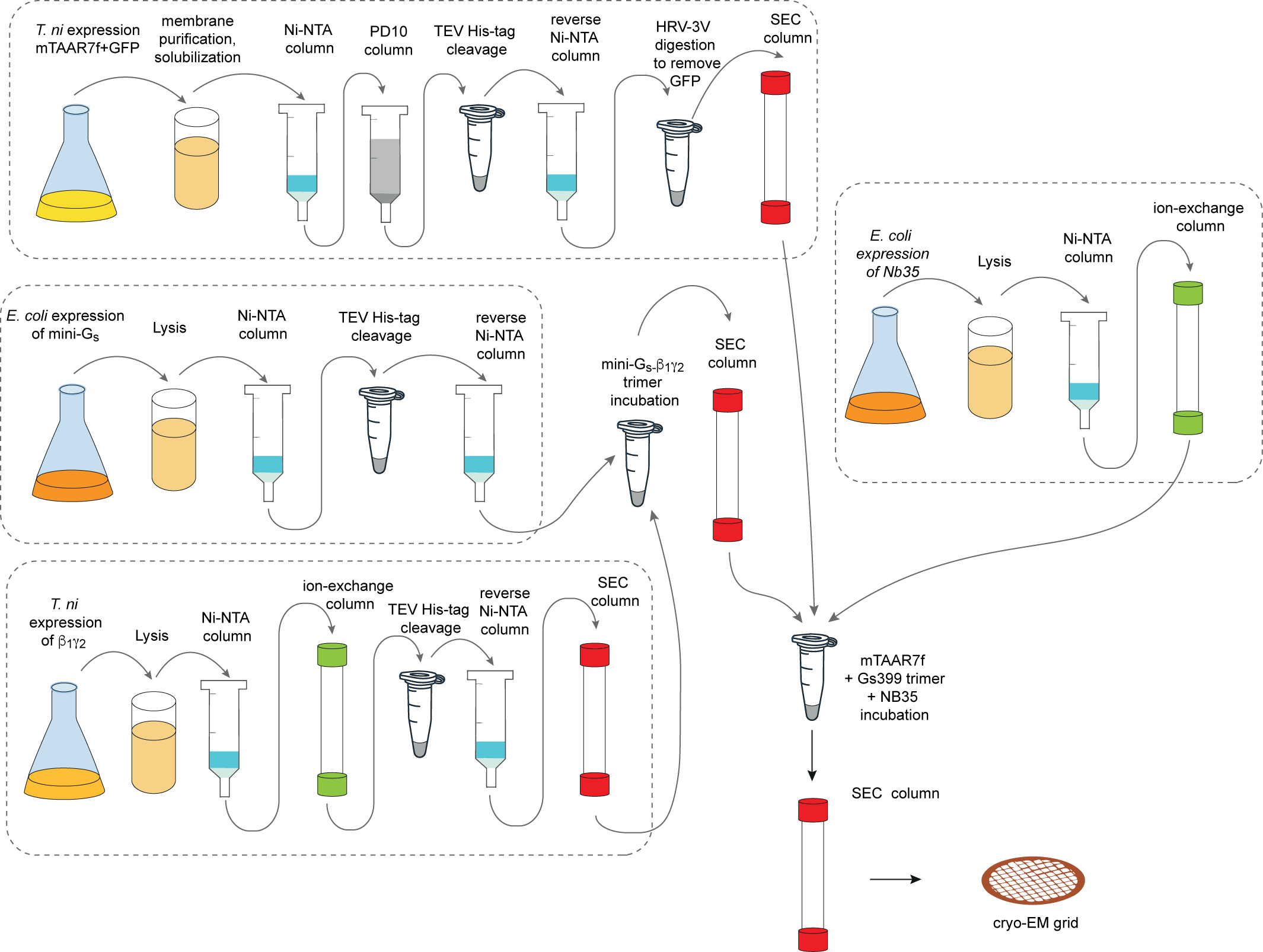
mTAAR7f purification scheme. Purification scheme for the preparation of the mTAAR7f–miniG_s_–Nb35 complex for structure determination by cryo-EM.

**Extended Data Fig. 2.**
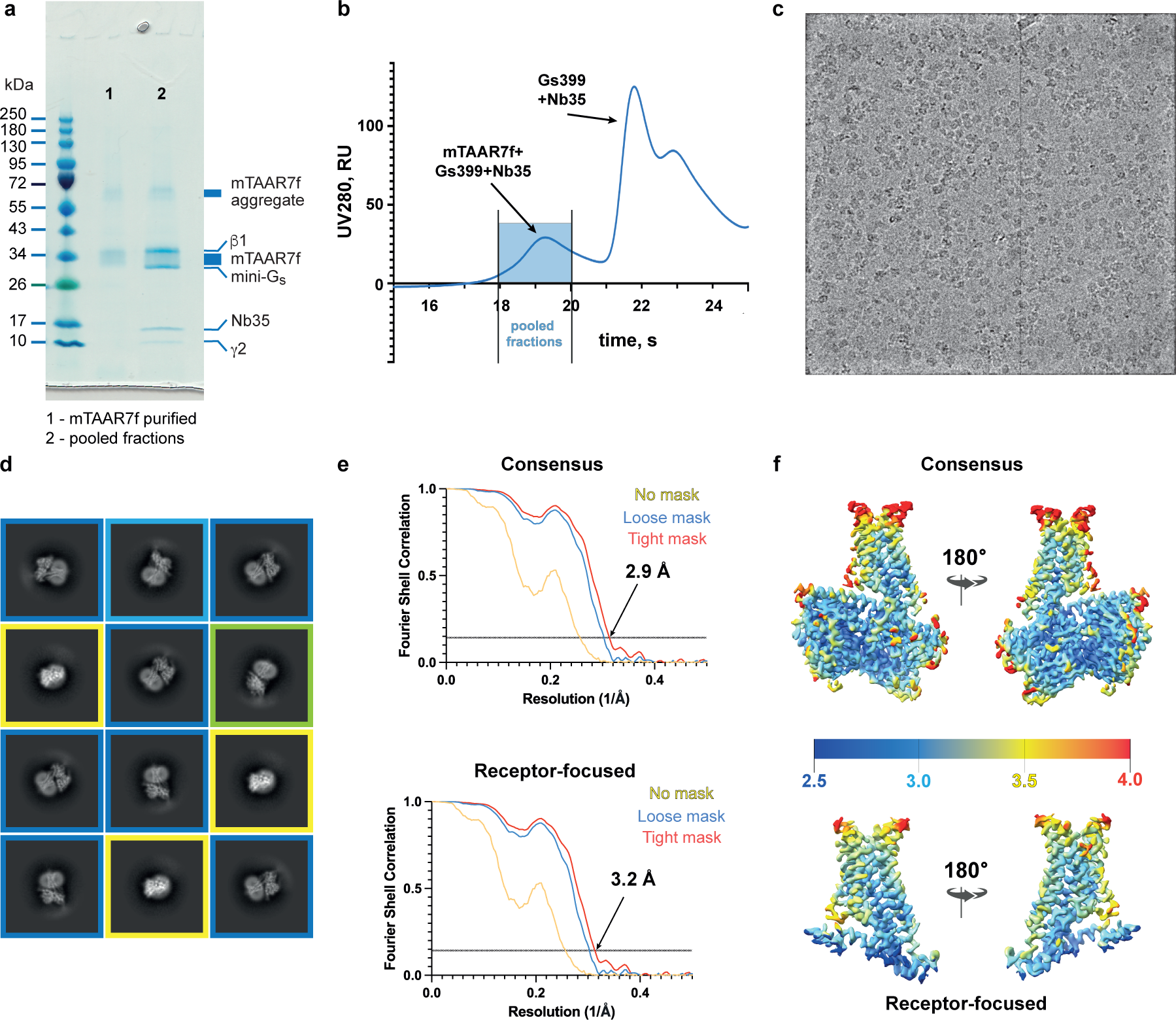
Cryo-EM of the mTAAR7f–mini-G_s_–Nb35 complex and single-particle reconstruction. **a**, Coomassie Blue-stained SDS-PAGE gel of purified mTAAR7f (lane 1) and pooled fractions of the mTAAR7f–miniG_s_–Nb35 complex after gel filtration (lane 2). Individual components are indicated. **b**, Gel filtration trace of the mixture of mTAAR7f with miniG_s_ and Nb35. **c**, A representative cryo-EM micrograph (defocus −2.4 μm) from the collected dataset. **d**, Representative 2D class averages of the mTAAR7f–miniG_s_–Nb35 complex determined using the initial set of particles following several rounds of 2D classification. Class averages corresponding to similar particle orientations are marked with the same coloured frames: blue, side views; green, partial side view; yellow, top views. **e**, FSC curves of the receptor-focused and consensus reconstructions show an overall resolution of 3.2 Å and 2.9 Å, respectively, using the gold standard FSC of 0.143. **f**, Local resolution estimation of the receptor-focused and consensus maps of the mTAAR7f–miniG_s_–Nb35 as calculated by CryoSparc.

**Extended Data Fig. 3.**
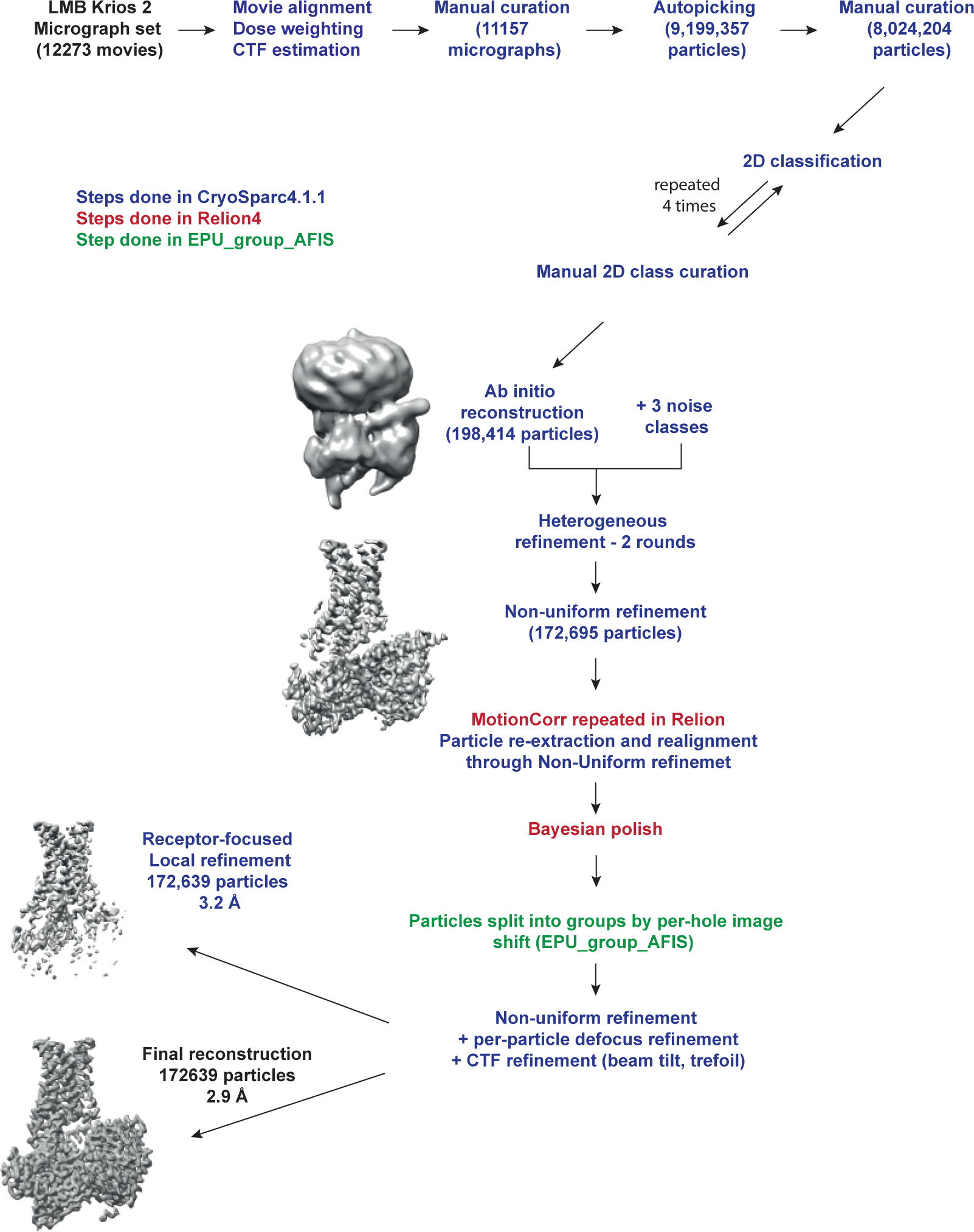
Flow chart of cryo-EM data processing. The dataset was collected in one session (48 h) on the LMB Krios 2 equipped with Falcon 4 detector. The movies were corrected for drift, beam-induced motion and radiation damage using CryoSparc motion correction implementation. After estimation of CTF parameters, the dataset was manually curated to exclude low quality micrographs. Particles were picked using a Gaussian blob and subjected to four rounds of 2D classification, after each round only species resembling a receptor-G protein complex were retained. Particles in the best 2D classes were subjected to two rounds of heterogenous refinement in CryoSparc versus three separately generated classes corresponding to picks without any structural features (noise classes). The output particles were subjected to one round of non-uniform refinement in CryoSparc resulting in a global resolution of 3.05 Å. To perform post-processing steps in Relion, motion correction was repeated in Relion followed by particle re-extraction and realignment. Bayesian polishing was performed in Relion, and particles were also split into AFIS groups using EPU_group_AFIS script. The final step of non-uniform refinement coupled to per-particle defocus refinement and per-particle CTF refinement, including beam-tilt, trefoil and tetrafoil corrections, was performed in CryoSparc. The final model based on 172,639 particles achieved a global resolution of 2.9 Å, while the receptor-focused map achieved a global resolution of 3.2 Å. Resolution of the models after refinements was calculated with the gold-standard FSC of 0.143 in CryoSparc (Extended Data Fig. 2).

**Extended Data Fig. 4.**
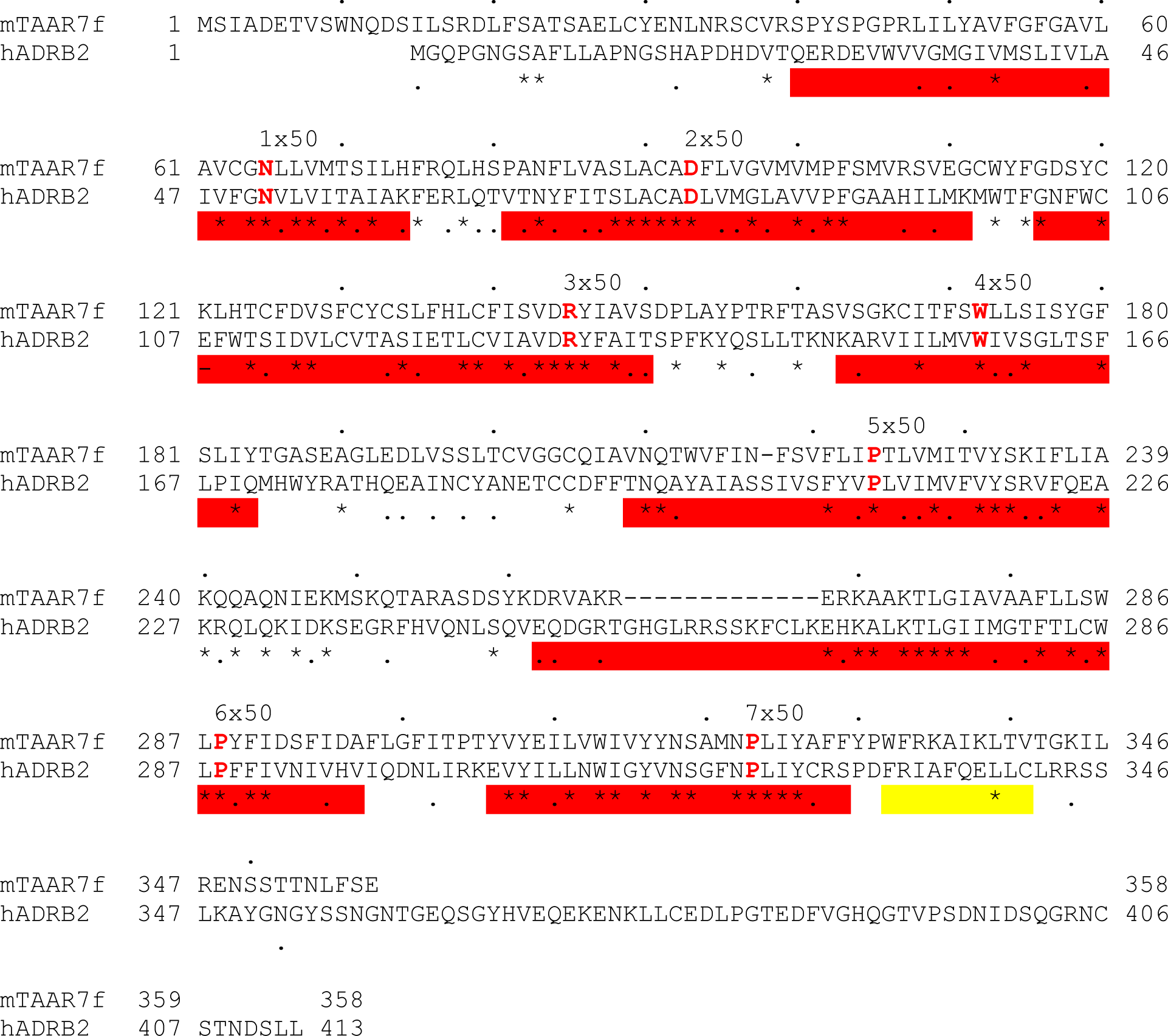
Alignment of the amino acid sequences of mTAAR7f and β_2_AR. Red bars, transmembrane regions; yellow bar, amphipathic helix 8; red residues, Ballesteros Weinstein numbering system xx.50. The alignment was performed using the program MacVector.

**Extended Data Fig. 5.**
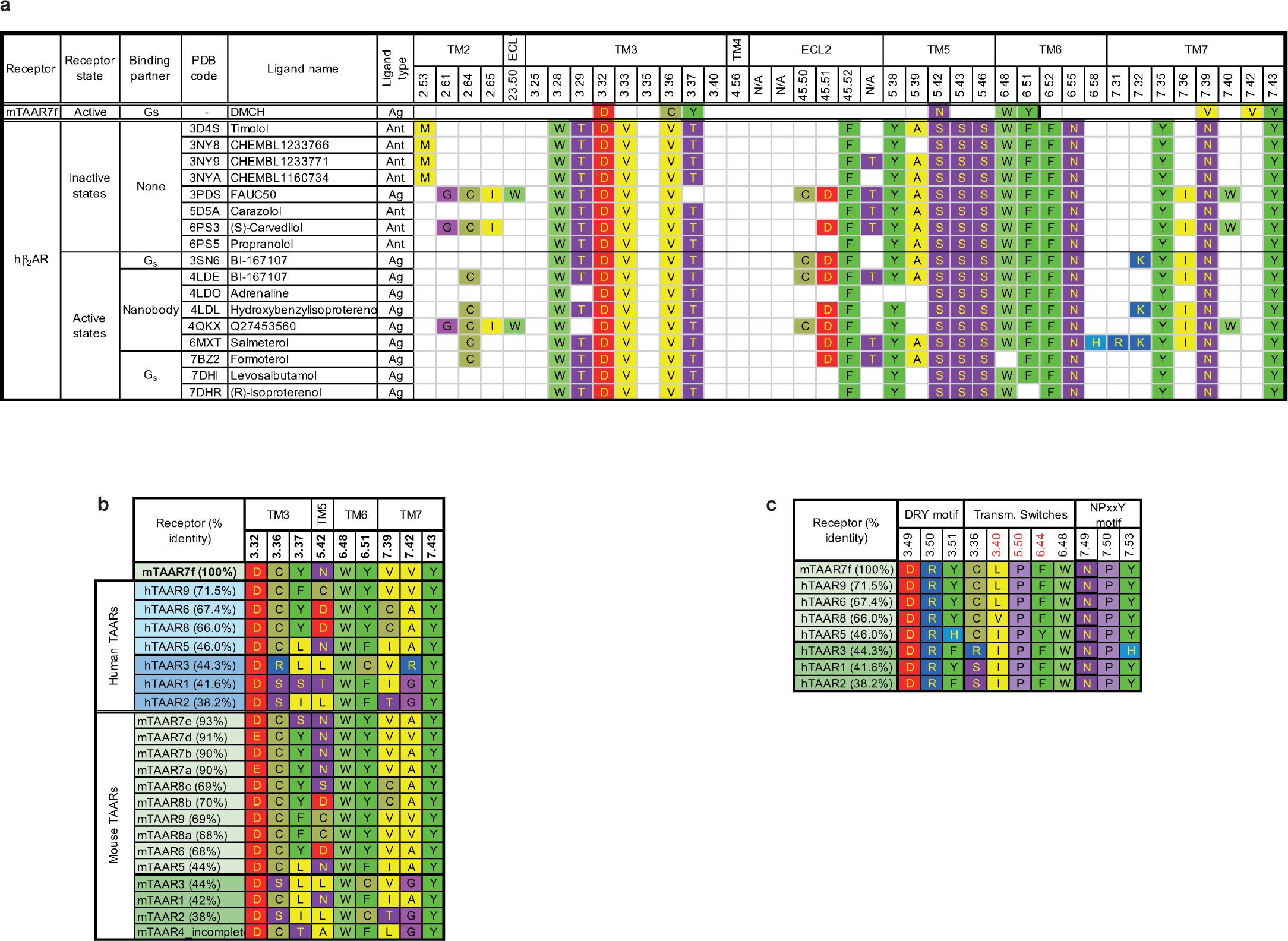
Sequence conservation in TAARs and the OBS. **a**, Amino acid residues within 3.9 Å of ligands in the mTAAR7f structure and structures of human β_2_AR. **b**, Amino acid residues within 3.9 Å of ligands in the mTAAR7f structure aligned with the equivalent residues in both human and mouse TAARs. Sequences were aligned using Clustal Omega^50,51^ and the percentage of the full-length receptor sequence identity to mTAAR7f was calculated using the web-based resource BLAST^52^. **c**, Conservation in TAARs of the D-R-Y motif, transmission switches (including the P-I-F motif, marked in red) and N-P-x-x-Y motifs. Sequences were aligned using Clustal Omega^50,51^. Percentage of the full-length receptor sequence identity to mTAAR7f was calculated using web-based resource BLAST^52^

**Extended Data Fig. 6.**
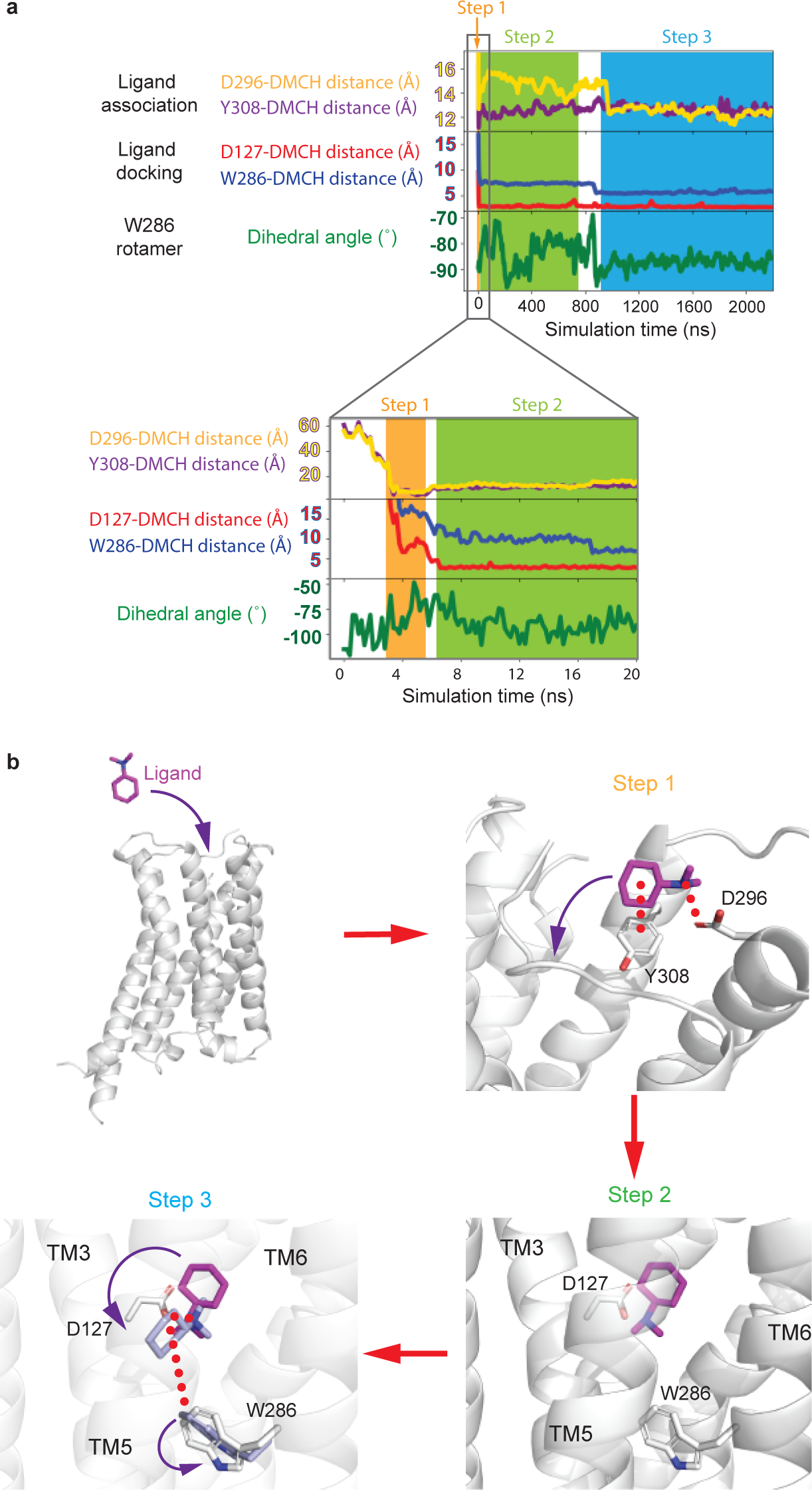
MD simulation of DMCH association to mTAAR7f. **a**, Four 2.2 μsec velocity MD simulation were performed on mTAAR7f (no G protein) in the presence of ligand outside the OBS. Three examples are shown in Extended Data Fig. 8 and one example is shown here where the ligand remained stably associated with the receptor at the end of the simulation. The ligand was observed to enter the OBS rapidly. Ligand clustering analysis identified specific residues (Asp296 and Tyr308) that associated with DMCH upon initial association with the receptor. The process is plotted visually through measuring distances between DMCH and the residues in the extracellular region (Asp296^6.58^ and Tyr308^7.35^) and in the OBS (Asp127^3.32^ and Trp286^6.48^). The motion of Trp286 is monitored through the variation in its Chi2 angle. **b**, Three step model for the binding of DMCH into mTAAR7f. Note that this simulation was performed on mTAAR7f in an active state and might not represent fully the trajectory in an inactive state in the absence of a G protein. However, given our understanding of the role of the G protein in closing the entrance of the OBS and decreasing its volume upon G protein coupling in the βARs, then the data here may represent an underestimate of the rate of ligand association.

**Extended Data Fig. 7.**
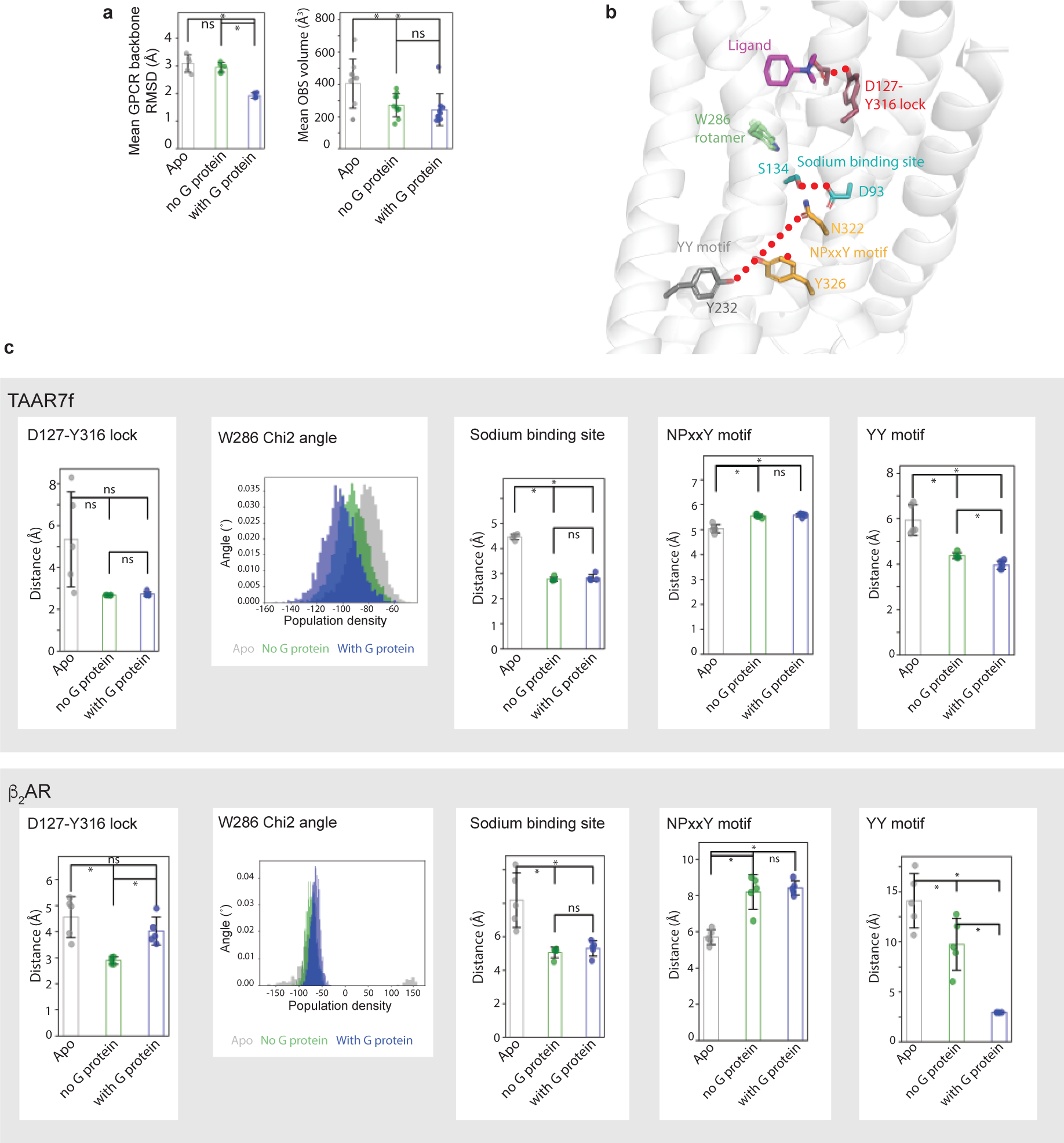
MD simulations of mTAAR7f and analysis of changes in activation switches. **a**, Five independent MD simulations were performed either on the mTAAR7f-mini-G_s_-DMCH complex (with G protein), on mTAAR7f-DMCH (no G protein) or on mTAAR7f alone (Apo). The mean GPCR backbone RMSD and the mean volume of the OBS are plotted for each simulation and found to increase significantly in the Apo simulations compared to when G protein and ligand are bound. **b**, Position of the transmission elements in mTAAR7f that were analysed to assess whether the receptor was remaining in the state defined by the cryo-EM structure. These included all the canonical transmission switches in Class A GPCRs. **c**, For each of the above simulations distances were plotted between residues that define the state of the transmission switches. No significant differences were observed in the OBS (D127-Y316 lock), but increases in distances were observed for both mTAAR7f and β_2_AR in the YY motif and the sodium, binding site, consistent with a tendency towards a more inactive state. No change was observed in the NPxxY motif in mTAAR7f, but β_2_AR changed towards a more inactive state. Changes in Chi2 angle of Trp286 in mTAAR7f show a tendency towards a more inactive state in the Apo simulation, but this is not in evidence in β_2_AR. Data for simulations on β_2_AR were obtained from GPCRmd. The error bars represent the SD and a t-test showed either no statistical difference (ns) or a statistical difference (*, p< 0.05) between data.

**Extended Data Fig. 8.**
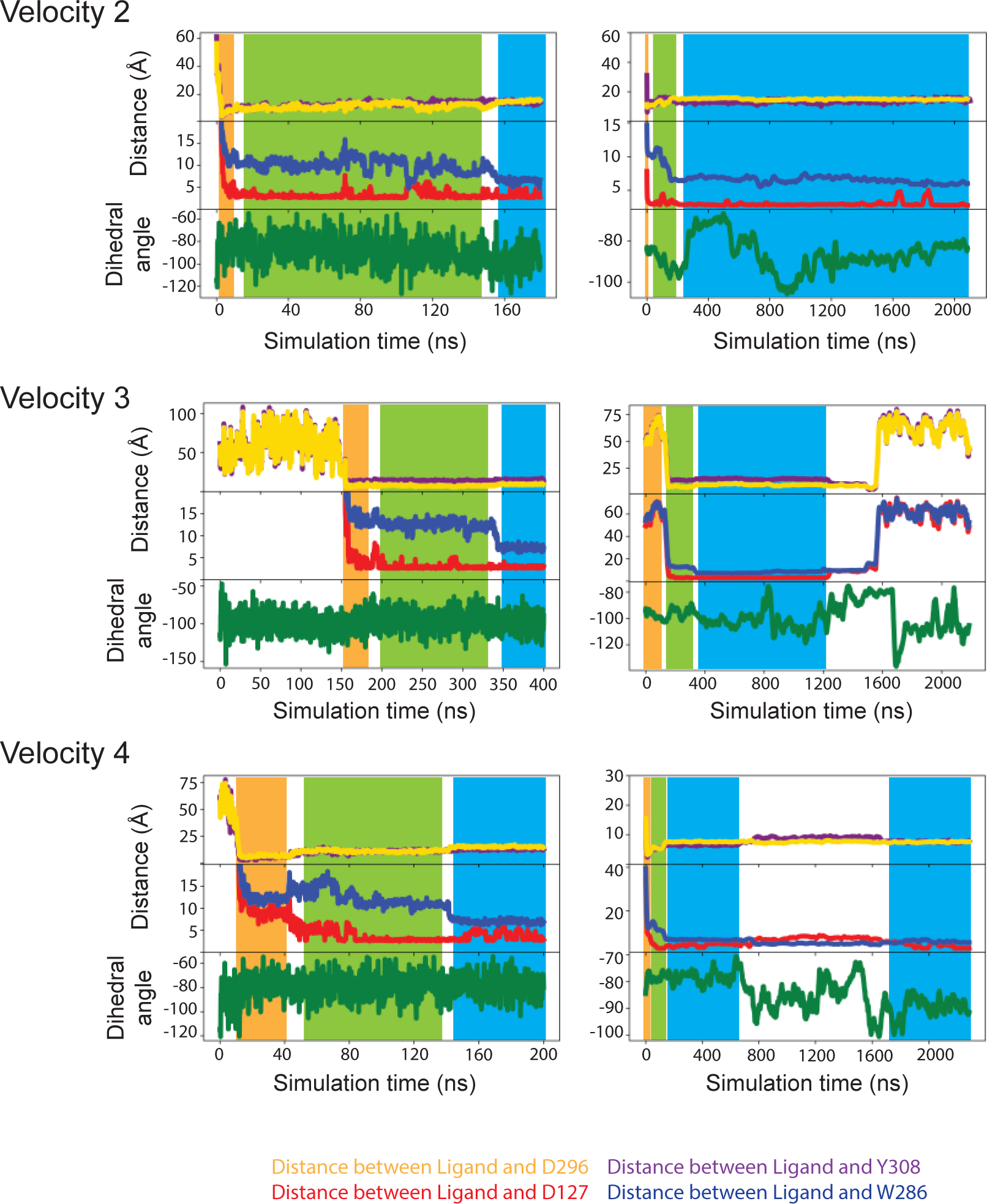
MD simulation of DMCH association to mTAAR7f. Three additional 2.2 μsec velocity MD simulation are shown of mTAAR7f (no G protein) in the presence of ligand outside the OBS. The colour scheme is identical to that in Extended Data Fig. 6a. The blue area in the traces (Step3) represents where the simulated position of DMCH is similar to that in the cryo-EM structure. In Velocity 2 the ligand is stable in Step 3, but in Velocity 3 the ligand dissociates and does not re-bind. In Velocity 4 the initially adopts a pose similar to the cryo-EM structure, but then rotates away form it, until re-adopting the cryo-EM pose 1 μsec later. Panels on the left represent the first 200-400 nsec and the panels on the right show the whole 2.2 μsec simulation.

**Extended Data Fig. 9.**
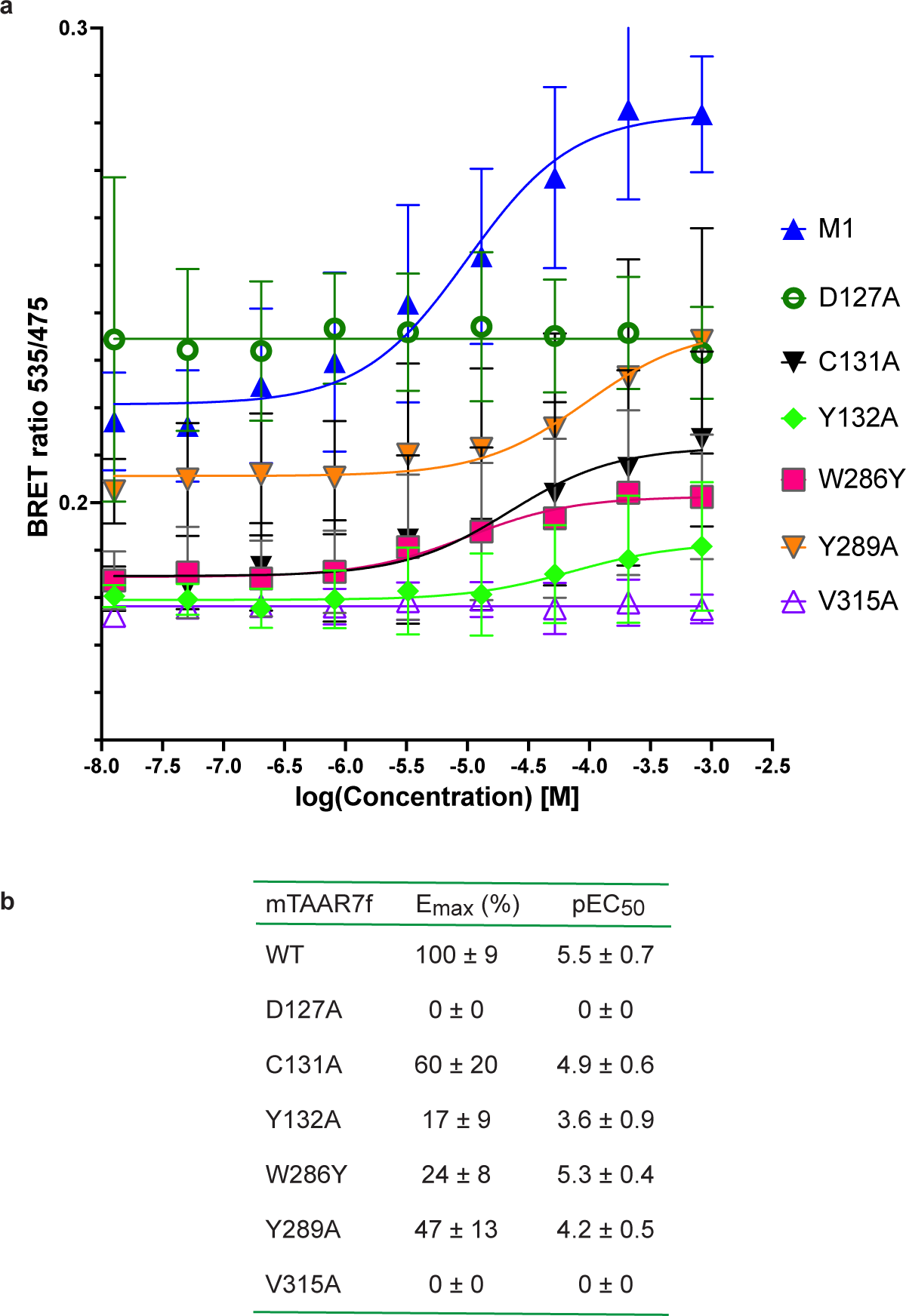
BRET data for G protein recruitment to mTAAR7f. **a**, BRET ratios measured for increasing concentrations of the agonist DMCH for the wild-type mTAAR7f (M1) and six mutants. **b**, Values for Emax and pEC50 determined from the data in panel **a**, with errors given as SEM. Three experiments were performed independently with single measurements per experiment.

